# Fluid restrictive resuscitation with high molecular weight hyaluronan infusion in early peritonitis-sepsis

**DOI:** 10.1101/2023.04.14.536966

**Authors:** Annelie Barrueta Tenhunen, Jaap van der Heijden, Paul Skorup, Marco Maccarana, Anders Larsson, Anders Larsson, Gaetano Perchiazzi, Jyrki Tenhunen

## Abstract

Sepsis is a condition with high morbidity and mortality. Prompt recognition and initiation of treatment is essential. Despite forming an integral part of sepsis management, fluid resuscitation may also lead to volume overload, which in turn is associated with increased mortality. The optimal fluid strategy in sepsis resuscitation is yet to be defined. Hyaluronan, an endogenous glycosaminoglycan with high affinity to water is an important constituent of the endothelial glycocalyx. We hypothesized that exogenously administered hyaluronan would contribute to endothelial glycocalyx integrity and counteract intravascular volume depletion in a fluid restrictive model of peritonitis. In a prospective, blinded model of porcine peritonitis-sepsis, we randomized animals to intervention with hyaluronan (n=8) or 0.9% saline (n=8). The animals received an infusion of 0.1% hyaluronan 6 ml/kg/h, or the same volume of saline, during the first two hours of peritonitis. Stroke volume variation and hemoconcentration were comparable in the two groups throughout the experiment. Cardiac output (p=0.008 and p=0.017) and diastolic blood pressure (p=0.015 and p=0.019) were higher in the intervention group during the infusion of hyaluronan, but these effects disappeared as the experiment proceeded. The increase in lactate was more pronounced in the intervention group (p=0.041) throughout the experiment, while concentrations of surrogate markers of glycocalyx damage; syndecan 1 (0.6 ± 0.2 ng/ml vs 0.5 ± 0.2 ng/ml, p=0.292), heparan sulphate (1.23 ± 0.2 vs 1.4 ± 0.3 ng/ml, p=0.211) and vascular adhesion protein 1 (7.0 ± 4.1 vs 8.2 ± 2.3 ng/ml, p=0.492) were comparable in the two groups at the end of the experiment. In conclusion, hyaluronan did not counteract intravascular volume depletion in early peritonitis sepsis. The intervention was associated with higher cardiac output and diastolic blood pressure, than placebo, during the infusion. However, the increase in lactate throughout the experiment was more pronounced in the intervention group.

## Introduction

Sepsis is a condition with high mortality in which a dysregulated host response to infection causes organ dysfunction [1]. While recovery depends on adequate anti-infective measures and therapy, the multifactorial nature of cardiovascular instability [2,3] is antagonized with fluids and vasopressors/inotropes [4]. Early fluid resuscitation is crucial to reverse the deleterious effects of tissue hypo-perfusion, but excessive fluid therapy is associated with increased mortality [5,6].

Microcirculatory perfusion disturbances are common in sepsis [7,8,9]. The deranged microcirculation is partly explained by endothelial dysfunction and glycocalyx degradation [9,10,11,12]. Endothelial dysfunction and glycocalyx degradation may lead to edema formation [13]. High molecular weight hyaluronan, HMW-HA (MW > 1000 kDa), is a highly hydrophilic constituent of the endothelial glycocalyx layer [14,15] and contributes to vascular integrity [11,16].

In states of inflammation, HMW-HA is degraded by several mechanisms [17,18,19] with concomitant shedding of the endothelial glycocalyx layer [20]. Exogenously administered glycosaminoglycans (HA, chondroitin sulphate) can restore shedded glycocalyx [16] and pericellular matrix [21] after hyaluronidase treatment. HA has been safely administered intravenously in humans [22] and reduces inflammation and lung injury in experimental sepsis [23].

In a previous study, we tested the effect of exogenously administered HMW-HA in peritonitis sepsis as adjuvant treatment to crystalloid fluid resuscitation. A post hoc analysis demonstrated a lower modified shock index (MSI=HR/MAP) during sepsis-peritonitis in the intervention group [24]. Crystalloid infusion per se increases plasma concentration of HA [25]. Therefore, in the present study we reduced the administered volume of crystalloid, with the hypothesis that exogenously administered HMW-HA contributes to endothelial glycocalyx integrity in sepsis and thereby counteracts intravascular volume depletion in a fluid restrictive model.

## Materials and methods

### Animals and ethic statements

The study was approved by the Animal Ethics Committee in Uppsala, Sweden (decision 5.8.18-01054/2017, DOUU 2019-014). The animals were cared for in strict accordance with the National Institute of Health guide for the care and use of Laboratory animals [26]. All efforts were made to minimize distress. After premedication, the animals received continuous intravenous anaesthesia and analgesia. The study was performed at the Hedenstierna Laboratory, Uppsala University, Sweden.

### Anaesthesia and instrumentation

Twenty male pigs (Sus scrofa domesticus) of mixed Swedish Hampshire and Yorkshire breeds (mean weight 30.4 ± 1.8 kg) received premedication with Zoletil Forte® (tiletamine and zolazepam) 6 mg/kg and Rompun® (xylazine) 2.2 mg/kg i.m. The animals were placed in supine position after adequate sedation was obtained. A peripheral intravenous catheter was placed in an auricular vein and a bolus of fentanyl of 5-10 µg/kg administered i.v. Anaesthesia was then maintained with ketamine 30 mg/kg/h, midazolam 0.1-0.4 mg/kg/h and fentanyl 4 µg/kg/h, in glucose 2.5% during the experiment.

Rocuronium 2.5 mg/kg/h was added as muscle relaxant after adequate depth of anaesthesia was assured by absence of reaction to pain stimulus between the front hooves. Ringer’s acetate was infused i.v. at a rate of 10 ml/kg/h during the first 30 minutes of the protocol. The animals were under deep anaesthesia during the whole experiment (six hours of peritonitis/sepsis), including during euthanasia (100 mmol KCl i.v.). Bolus doses of 100 µg fentanyl i.v. were administered if signs of distress or reaction to pain stimulus were noted.

The airway of the animals was secured via tracheostomy. A tube of an internal diameter of eight mm (Mallinckrodt Medical, Athlone, Ireland) was inserted in the trachea. Thereafter, volume controlled ventilation (Servo I, Maquet, Solna, Sweden) was maintained as follows: respiratory rate (RR) 25/min, tidal volume (V_T_) 8 ml/kg, positive end-expiratory pressure (PEEP) 8 cmH_2_O and inspired oxygen concentration (F_I_O_2_) 0.3. The settings of VT and PEEP were maintained throughout the protocol, while RR was adjusted aiming at a PaCO_2_ <6.5 kPa, and F_I_O_2_ was adjusted to keep PaO_2_ >10 kPa.

A pulmonary artery catheter for measurement of pulmonary artery pressures and cardiac output (CO) and a triple lumen central venous catheter for fluid infusions were inserted via the right jugular vein. An arterial catheter was inserted via the right carotid artery for blood pressure measurement and blood sampling. A PiCCO (Pulse index continuous cardiac output) catheter (Pulsion, Munich, Germany) was inserted via the right femoral artery for estimation of stroke volume variation (SVV) and extravascular lung water (EVLW). Blood gases were analysed immediately after sampling and executed on an ABL 3 analyser (Radiometer, Copenhagen, Denmark). Hemoglobin (hgb) and hemoglobin oxygen saturation were analysed with a hemoximeter OSM 3 calibrated for porcine hemoglobin (Radiometer, Copenhagen, Denmark).

A midline laparotomy was performed. The bladder was catheterized for urinary drainage and an incision was made in the caecum, feces were collected and thereafter the incision in the cecal wall was closed. After insertion of a large-bore intra-peritoneal drain, the abdominal incision was closed.

### Study protocol

To yield a stock solution of 1% (10 mg/ml), five grams of HMW-HA 1560 kDa (Sodium hyaluronate Lot# 027362 HA15M-5, Lifecore Biomedical LCC, Chaska, MN, USA) was dissolved in 500 ml 0.9% saline. The solution of 1% HMW-HA was produced under sterile condition in laminar airflow and stored as 50 ml aliquots at −20◦C. On the day of experiment, aliquots were diluted 1:10 in 0.9% saline, to yield 0.1% concentration.

Peritonitis was induced, after baseline measurements, via a peritoneal instillation of autologous feces (2 g/kg body weight in 200 ml warmed 5% glucose solution), after which the abdominal wall was closed.

### Experimental design

The experimental time line is presented in figure 1. The animals were randomized in two steps (block randomization, sealed opaque envelope), first to peritonitis (n=16) or time control (n=4), then into two treatment groups: intervention with HMW-HA (n=8) or control group (n=8). The study was prospective and the researchers were blinded for the group allocation until a master file for the whole experiment was produced.

**Figure 1.**
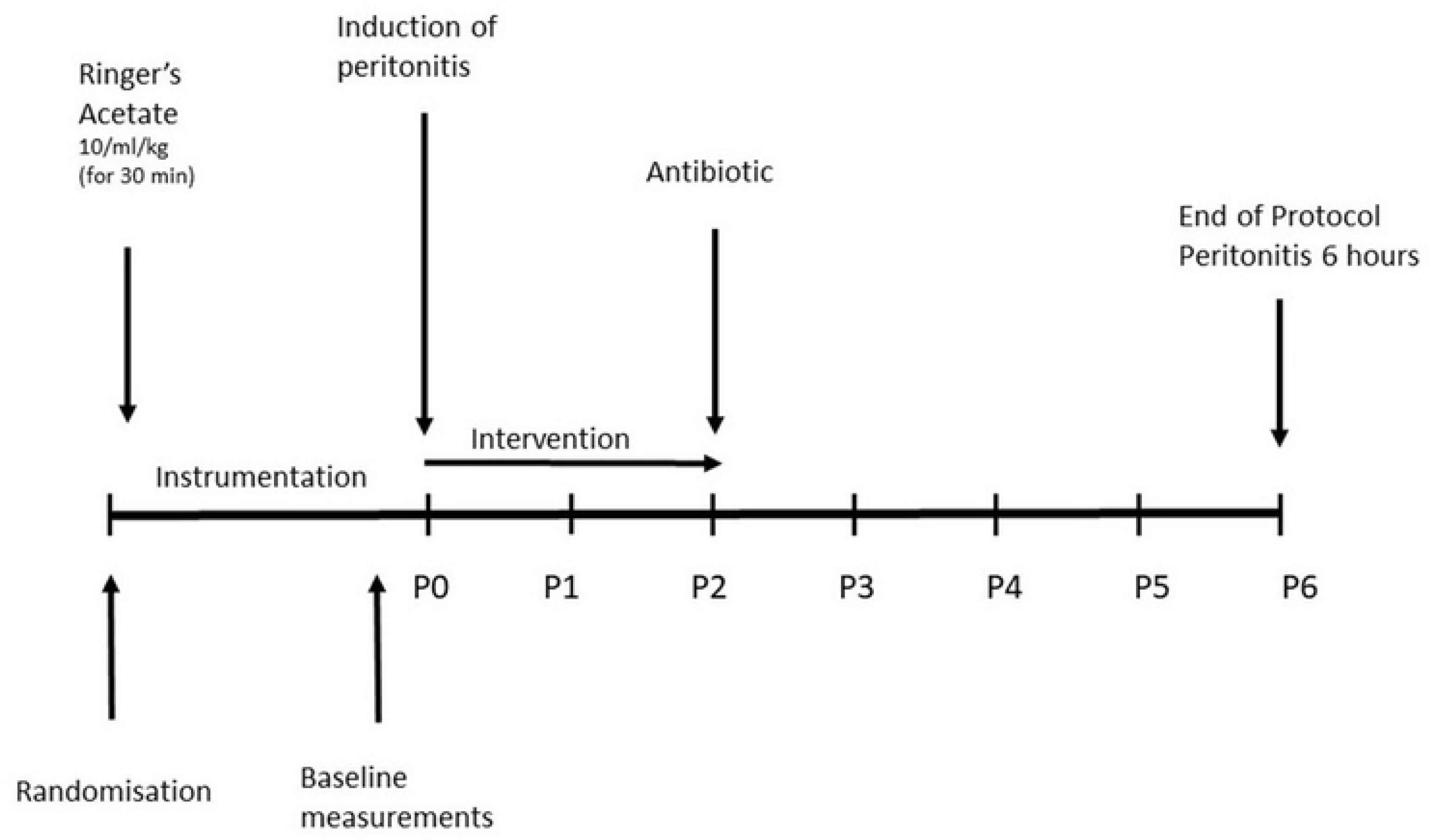
Experimental timeline. P0 is the induction of peritonitis, followed by P1 (one hour after peritonitis induction), P2, P3, P4, P5 and P6 (six hours after peritonitis induction and end of protocol).

The intervention was started at the time of the laparotomy with 0.1 % solution HMW-HA, administered with a rate of 6 mg/kg/hour (6 ml/kg/h) for two hours. The control group received the same volume of vehicle (0.9% saline, 6 ml/kg/h) as an infusion over two hours. After two hours duration of peritonitis two grams of Piperacillin/Tazobactam in 10 ml of 0.9 % saline was administered i.v. If the animals developed circulatory instability (defined as MAP < 55 mmHg > five minutes) an infusion of Norepinephrine (40 µg/ml) was started with the rate of 5 ml/hour and increased stepwise, aiming at maintaining a MAP > 55 mmHg. No additional fluids were administered.

### Analyses and physiologic parameters

The primary endpoint parameter, SVV, was measured at baseline and every hour for the following six hours duration of the experiment, simultaneously with EVLW and arterial blood gas analysis (time points P1-P6). Concomitantly, hemodynamic parameters (systemic arterial and pulmonary arterial pressures, CO, heart rate), respiratory parameters (F_I_O_2_, SaO_2_, ETCO_2_, peak pressure, plateau pressure, dynamic and static compliance) and urine output were measured. Mixed venous blood gas analysis, collection of plasma samples and arterial blood for bacterial cultures were drawn at baseline and at peritonitis duration of one, two, three and six hours.

### Cytokine and HA analyses, VAP1, Syndecan 1, Heparan Sulphate

Porcine-specific sandwich ELISAs were used for the determination of TNF-α and interleukin-6 (IL-6) in plasma (DY690B (TNF-α), DY686 (IL-6), R&D Systems, Minneapolis, MN, USA). The ELISAs had total coefficient of variations (CV) of approximately 6%. A commercial ELISA kit (Hyaluronan DuoSet, DY3614, R&D Systems, Minneapolis, MN, USA) was used to measure the hyaluronan concentration. Porcine specific ELISA kits were used to analyze plasma concentration of vascular adhesion protein 1 (VAP 1) (MyBioSource cat. No. MBS9364679), syndecan 1 (MyBioSource cat. No. MBS2703970) and heparan sulphate (MyBioSource cat. No. MBS265068).

### Total protein and osmolality

Total protein was analysed using a Mindray BS380 (Mindray, Shenzhen, China) with reagents from Abbott Laboratories (7D73-22, Abbott Park, IL, USA). Osmolality was measured using an OsmoPRO Multi-Sample Micro-Osmometer (I&L Biosystems, Königswinter, Germany).

### Bacterial investigations

From a sterile arterial catheter, 0.5 ml blood was drawn for quantitative blood cultures. 100 µl was cultured on three separate cysteine lactose electrolyte deficient (CLED) agar plates and cultured at 37°C overnight. Colony forming units (CFU) were quantified with viable count technique the following day. The median of counted CFU/mL was calculated. CFU on one of three CLED plates from a time point was interpreted as a contamination. More than 1 CFU/mL were considered a positive blood culture.

### Statistical analysis

To determine sample size we used data from a previous peritonitis protocol where the stroke volume variation was used as a guide to fluid therapy. The control group had a standard deviation of ± 2 % at baseline. Aiming at detecting a difference of 3 per cent units of SVV between groups, a power of 0.8 and a significance level of < 0.05 justified a sample size of eight animals in each group.

The Shapiro-Wilk’s test was used to test data for normality. The two-tailed Student’s t-test, the Mann-Whitney U test and the two-way ANOVA were used to compare the two groups, pending distribution of data.

Data are expressed as mean ± SD or median (IQR) according to distribution of data. We conducted the statistical analyses using SPSS v. 28.0.0 software (SPSS, Inc., Chicago, IL, USA). A p-value of < 0.05 was considered statistically significant.

## Results

All animals survived to the end of the experiment. In the intervention group as well as in the control group six of eight animals presented with circulatory instability (defined as MAP < 55 mmHg > five minutes) within the time frame of the protocol.

### Hyaluronan plasma concentration

The hyaluronan concentration was comparable in the two groups with 67 ± 14 ng/ml in intervention group and 85 ± 25 ng/ml in control group (p = 0.103) at baseline. In the control group no statistically significant dynamics in the hyaluronan concentration was detected (one-way ANOVA, p = 0.580). In the intervention group the hyaluronan concentration peaked after the two-hour infusion with 158708 ± 57242 ng/ml and declined to 57801 ± 32153 ng/ml at 6 hours (p = 0.002) (Figure 2).

**Figure 2.**
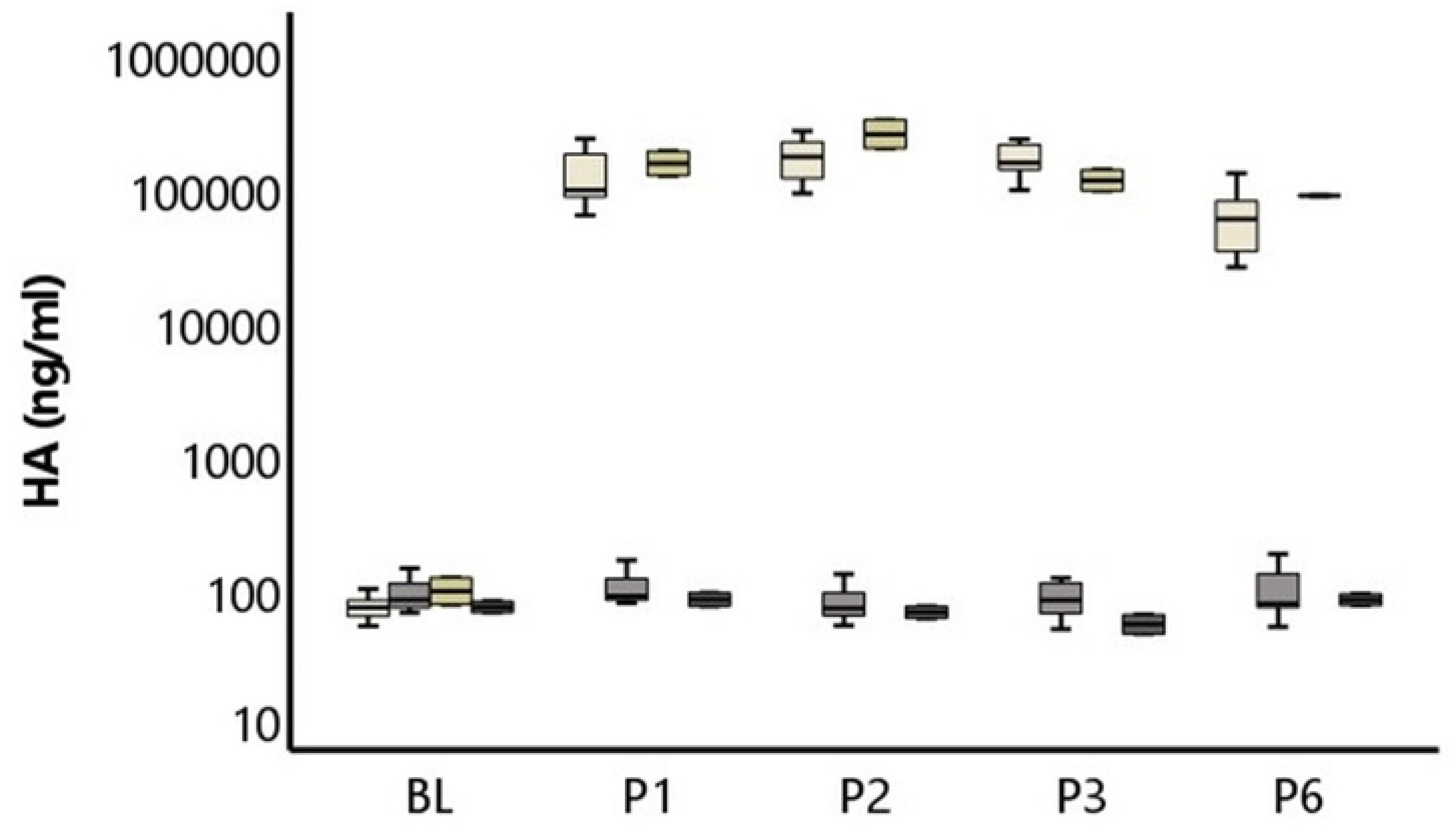
Hyaluronan concentration (HA) in plasma in the different groups at baseline (BL), and after one (P1), two (P2), three (P3) and six (P6) hours of peritonitis. White represent intervention and grey control group, whereas yellow and dark grey represent time control animals (HA vs control).

### Hemodynamics

Mean time to onset of circulatory instability in the intervention group was 4.4 ± 0.9 and 4.7 ± 0.3 h (p = 0.417) in the control group. MAP declined in both groups during the experiment and MSI, heart rate, lactate and temperature increased equally in the two groups. This was associated with, and preceded by, an increase in SVV and haemoglobin, comparable in both groups.

Diastolic blood pressure decreased comparably in the two groups as a function of time (p = 0.313). However, at P2 (p = 0.015) and P3 (p = 0.019), diastolic blood pressure was higher in the intervention group (see Table 1 for all hemodynamic parameters).

**Table 1.** Hemodynamics.

|  | Group | Baseline<br>(n=8+8) | 1 hour of<br>peritonitis<br>(n=8+8) | 2 hours of<br>peritonitis<br>(n=8+8) | 3 hours of<br>peritonitis<br>(n=8+8) | 6 hours of<br>peritonitis<br>(n=8+8) | p value |
| --- | --- | --- | --- | --- | --- | --- | --- |
| BE<br>(mEq/l) | HA | 2.0 ± 3.9 | 2.1 ± 2.3 | 1.5 ± 1.7 | 0.5 ± 1.5 | -4.1<br>± 3.3 | 0.443 |
|  | Control | 4.6 ± 1.5 | 1.8 ± 2.0 | 1.7 ± 1.9 | 1.4 ± 2.0 | -1.2 ± 3.9 |  |
| CI<br>(l/min/kg) | HA | 0.09 ± 0.02 | 0.11 ± 0.01 | 0.10 ± 0.01 | 0.10 ± 0.01 | 0.07 ± 0.02 | 0.142 |
|  | Control | 0.09 ± 0.02 | 0.09 ± 0.01 | 0.09 ± 0.01 | 0.09 ± 0.01 | 0.08 ± 0.01 |  |
| CO<br>(l/min) | HA | 2.8 ± 0.7 | 3.2 ± 0.5 | 3.2 ± 0.4 | 3.0 ± 0.4 | 2.1 ± 0.5 | 0.104 |
|  | Control | 2.5 ± 0.6 | 2.7 ± 0.2 | 2.7 ± 0.3 | 2.8 ± 0.4 | 2.3 ± 0.3 |  |
| CVP<br>(mmHg) | HA | 9 ± 2 | 8 ± 3 | 8 ± 3 | 7 ± 3 | 7 ± 2 | 0.920 |
|  | Control | 9 ± 3 | 9 ± 3 | 9 ± 3 | 8 ± 3 | 7 ± 3 |  |
| DBP<br>(mmHg) | HA | 71 ± 10 | 66 ± 10 | 69 ± 7 | 67 ± 9 | 42 ± 13 | 0.313 |
|  | Control | 75 ± 21 | 59 ± 10 | 58 ± 8 | 58 ± 5 | 45 ± 10 |  |
| ERO2 | HA | 0.6 ± 0.2 | 0.5 ± 0.1 | 0.5 ± 0.1 | 0.5 ± 0.1 | 0.7 ± 0.1 | 0.179 |
|  | Control | 0.5 ± 0.0 | 0.5 ± 0.1 | 0.5 ± 0.1 | 0.5 ± 0.1 | 0.6 ± 0.0 |  |
| Hb<br>(g/l) | HA | 89 ± 10 | 99 ± 8 | 110 ± 9 | 120 ± 10 | 127 ± 8 | 0.794 |
|  | Control | 93 ± 6 | 100 ± 10 | 106 ± 13 | 114 ± 13 | 129 ± 13 |  |
| HR<br>(BPM) | HA | 94 ± 9 | 102 ± 21 | 129 ± 41 | 159 ± 51 | 178 ± 41 | 0.203 |
|  | Control | 94 ± 19 | 86 ± 14 | 96 ± 23 | 108 ± 26 | 170 ± 44 |  |
| Lactate | HA | 1.2 ± 0.3 | 2.0 ± 0.5 | 1.9 ± 0.6 | 1.9 ± 0.7 | 3.2 ± 1.0 | 0.041* |
| (mmol/l) | Control | 0.9 ± 0.4 | 2.0 ± 0.6 | 2.0 ± 0.9 | 1.8 ± 1.0 | 1.7 ± 0.7 |  |
| MAP | HA | 89 ± 10 | 79 ± 13 | 81 ± 7 | 79 ± 10 | 52 ± 12 | 0.668 |
| (mmHg) | Control | 88 ± 19 | 73 ± 11 | 71 ± 9 | 71 ± 5 | 54 ± 12 |  |
| MSI | HA | 1.1 ± 0.2 | 1.3 ± 0.3 | 1.6 ± 0.4 | 2.1 ± 0.9 | 3.8 ± 1.8 | 0.725 |
|  | Control | 1.1 ± 0.4 | 1.2 ± 0.2 | 1.4 ± 0.3 | 1.5 ± 0.4 | 3.5 ± 1.8 |  |
| pH | HA | 7.40 ± 0.06 | 7.37 ± 0.04 | 7.35 ± 0.02 | 7.35 ± 0.04 | 7.31 ± 0.05 | 0.941 |
|  | Control | 7.43 ± 0.02 | 7.37 ± 0.02 | 7.38 ± 0.02 | 7.36 ± 0.03 | 7.33 ± 0.07 |  |
| SV | HA | 29 ± 6 | 33 ± 7 | 26 ± 8 | 21 ± 7 | 13 ± 4 | 0.413 |
| (ml) | Control | 28 ± 7 | 32 ± 3 | 29 ± 5 | 26 ± 4 | 14 ± 4 |  |
| SVI | HA | 1.0 ± 0.2 | 1.1 ± 0.2 | 0.9 ± 0.3 | 0.7 ± 0.2 | 0.4 ± 0.1 | 0.457 |
| (ml/kg) | Control | 0.9 ± 0.2 | 1.1 ± 0.1 | 1.0 ± 0.1 | 0.9 ± 0.1 | 0.5 ± 0.1 |  |
| SvO<sub>2</sub> | HA | 43 ± 13 | 48 ± 9 | 48 ± 8 | 46 ± 10 | 31 ± 13 | 0.404 |
| (%) | Control | 50 ± 4 | 45 ± 11 | 45 ± 9 | 51 ± 8 | 37 ± 4 |  |
| SVV | HA | 7 ± 2 | 9 ± 2 | 9 ± 4 | 10 ± 3 | 15 ± 6 | 0.636 |
| (%) | Control | 9 ± 3 | 9 ± 3 | 10 ± 3 | 12 ± 5 | 17 ± 6 |  |
| T | HA | 40.2 ± 0.8 | 40.2 ± 0.8 | 40.6 ± 1.0 | 41.1 ± 1.1 | 41.3 ± 1.3 | 0.928 |
| (° C) | Control | 39.7 ± 0.5 | 39.9 ± 0.8 | 40.2 ± 0.9 | 40.7 ± 1.0 | 41.6 ± 0.8 |  |
| Wedge | HA | 12 ± 2 | 10 ± 3 | 10 ± 2 | 10 ± 2 | 12 ± 7 | 0.953 |
| Pressure | Control | 11 ± 2 | 11 ± 3 | 11 ± 3 | 10 ± 3 | 11 ± 3 |  |
| (mmHg) |  |  |  |  |  |  |  |

Table 1. BE (base excess), CI (Cardiac index), CO (Cardiac output), CVP (central venous pressure), DBP (diastolic blood pressure), ERO_2_ (Oxygen extraction ratio), Hb (hemoglobin), HR (heart rate), BPM (beats per minute), MAP (mean arterial pressure), MSI (modified shock index, heart rate/MAP), SV (stroke volume), SVI (stroke volume index), SvO_2_ (mixed venous saturation), SVV (stroke volume variation), T (temperature). Values expressed as mean ± SD. Groups compared as a function of time (two way ANOVA), p value < 0.05 was considered to be statistically significant, in table marked with *.

There was no difference in CO between intervention and control group at baseline (p = 0.510). CO was higher in the intervention group at P1 (p = 0.008) and at P2 (p = 0.017). The difference disappeared at P3 (one hour after discontinued infusion) (p = 0.170). The difference applied also when correcting for Body Weight (CI: l/kg/min), with a higher CI in the intervention group at P1 (p = 0.013) and at P2 (p = 0.016), but at P3 the difference was no longer discernible (p = 0.247). When comparing SV and SVI at different time points, there was no statistically significant difference between groups. Heart rate was higher in the intervention group at P3 (p = 0.028), but this difference disappeared as the experiment proceeded.

Lactate increased in both groups from normal values at baseline, with a more pronounced increase in the intervention group during the experiment. Oxygen extraction ratio increased in both groups comparably and mixed venous oxygen saturation (SvO_2_) decreased without a discernible difference between groups throughout the experiment. pH and base excess (BE) were comparable in the two groups throughout the experiment, but BE was significantly lower in the intervention group at P5 (p = 0.021) and P6 (p = 0.028). The groups did not differ in central venous pressure (CVP) or wedge pressure as a function of time, values were normal throughout the protocol.

Norepinephrine requirement was comparable in the two groups 0.42 (0.97) µg/kg/min in the intervention group vs 0.37 (0.32) µg/kg/min in control group (p = 0.589) as well as the weight gain of 1.8 ± 0.4 kg vs 1.8 ± 0.4 kg. The time control animals were hemodynamically stable throughout the experiment (S1, Table).

### Respiratory parameters

Respiratory parameters were comparable between the two groups at baseline. PaO_2_/F_I_O_2_ ratio decreased in both groups, the decline reached statistical significance in the control group (one-way ANOVA, p = 0.02) but not in the intervention group (p = 0.182), there was no difference between groups as a function of time. Static compliance also decreased in the control group (one-way ANOVA p = 0.016), but not in intervention group (one-way ANOVA, p = 0.134), there was no difference between groups as a function of time. SaO_2_, peak pressure, plateau pressure, MPAP and EVLW were comparable in the two groups throughout the experiment (Table 2).

**Table 2.**
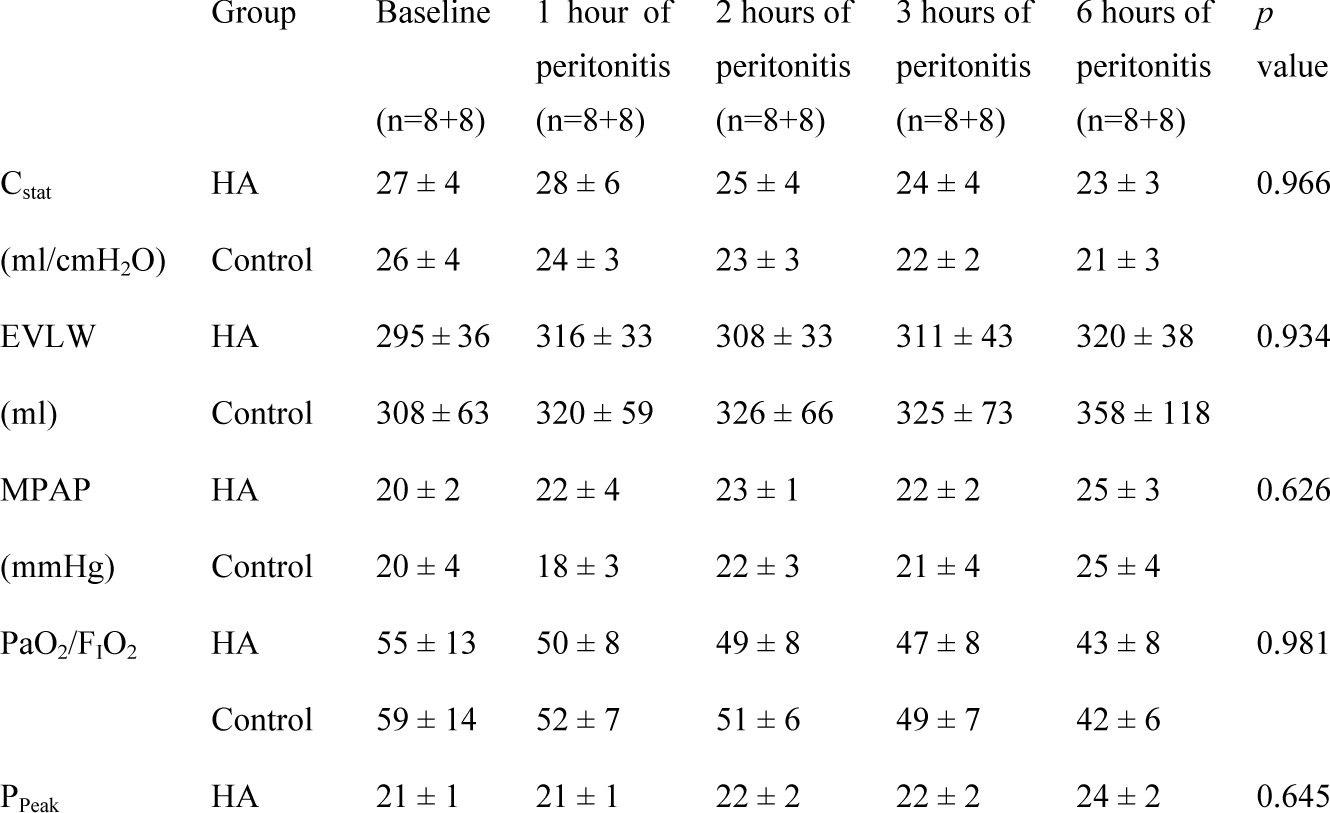
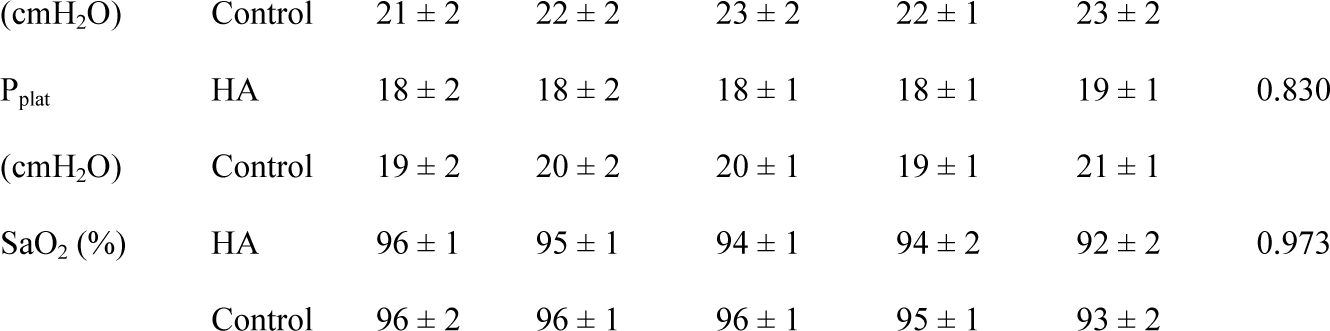
Respiratory parameters.

Table 2. Static compliance (C_stat_), extra vascular lung water (EVLW), mean pulmonary arterial pressure (MPAP), arterial oxygen partial pressure to fractional inspired oxygen ratio (PaO_2_/F_I_O_2_), peak pressure (P_Peak_), plateau pressure (P_plat_), arterial oxygen saturation (SaO_2_). Values expressed as mean ± SD. Groups compared as a function of time (two-way ANOVA).

### IL-6, TNFα and blood cultures

Plasma concentrations of IL-6 increased in both intervention and control groups from baseline throughout the experiment (six hours of peritonitis) from 80 (150) to 4316 (3940) pg/ml from 80 (0) to 4145 (2336) pg/ml, no difference between groups over time (p = 0.865). TNFα increased comparably in both groups (p = 0.950). Blood cultures were positive in three animals in each group during the observation period.

### Syndecan 1, heparan sulphate and VAP 1

Plasma concentration of Syndecan 1 was comparable in the two groups at six hours of peritonitis 0.6 ± 0.2 ng/ml vs 0.5 ± 0.2 ng/ml (p = 0.292). Heparan sulphate concentration did not differ between the two groups 1.23 ± 0.2 vs 1.4 ± 0.3 ng/ml (p = 0.211).

Plasma concentration of VAP1 was at six hours of peritonitis comparable between intervention and control groups, 7.0 ± 4.1 vs 8.2 ± 2.3 ng/ml (p = 0.492).

### Plasma protein and osmolality

Total protein was 40.7 (5.7) g/l in intervention group at baseline and 37.6 (3.5) g/l at six hours of peritonitis (p = 0.207), vs 39.6 (5.3) g/l at baseline in the control group and 35.8 (6.0) g/l at the end of the protocol. There was no difference between groups as a function of time (p = 0.703).

Osmolality was 284 (26) mOsm/kg at baseline in intervention group and 275 (29) mOsm/kg at six hours, vs 283 (15) mOsm/kg at baseline and 275 (8) mOsm/kg at six hours in the control group (p = 0.063). No difference between the groups throughout the experiment (p = 0.822).

## Discussion

In the present study, we opted for a restrictive fluid strategy in a porcine model of peritonitis sepsis. We hypothesized that the intervention with HMW-HA without additional crystalloid infusion would suffice to better maintain intravascular volume, blood circulation and the integrity of the glycocalyx. We chose SVV, a dynamic surrogate marker of intravascular volume as our primary end point parameter. Contrary to our hypothesis, HMW-HA administered in the early course of porcine peritonitis did not counteract the signs of intravascular hypovolemia as depicted by increasing SVV. At the same time, two-hour infusion of hyaluronan was associated with higher cardiac output/cardiac index and higher diastolic arterial pressure during and immediately following infusion. Hemoconcentration, increasing hgb/hematocrit (capillary leakage) and surrogate markers of endothelial glycocalyx damage (syndecan 1, heparan sulfate, VAP 1) did not differ between the two groups during the whole of the experiment. The inflammatory response as reflected by concentrations of selected cytokines in plasma was comparable in the two groups.

SVV is a validated method to assess preload responsiveness in mechanically ventilated, critically ill patients [27], with the magnitude of the cyclic changes in left ventricular stroke volume being proportional to volume responsiveness [28]. Apart from being a means of dynamic monitoring to guide fluid resuscitation, SVV can also be used to evaluate intravascular volume status [29]. In the present study no difference in SVV was observed between groups during the six hours of peritonitis sepsis, which was in accordance with the finding of comparable increase in plasma levels of hgb between the two groups; marker of hemoconcentration and indicative of loss of intravascular volume.

Plasma concentrations of HA peaked with a mean value of 158708 ± 57242 ng/ml, directly after the intervention was stopped (two hours of infusion), followed by a decline already at six hours to 57801 ± 32153 ng/ml. In a study by Hamilton et al. 2009 [22], the same dose (12 mg/kg as an infusion administered over two hours) of HA (mean MW 280 kDa) in healthy volunteers resulted in a two times higher peak concentration and the decline was not as pronounced at six hours.

Focusing on the period of intervention in the current study, some differences were statistically significant between the two groups after an hour-by-hour analyses. CO was higher in the intervention group during the infusion (P1 and P2). One hour after the infusion was paused (P3) CO was already comparable in the two groups. Diastolic blood pressure was significantly higher in the intervention group at P2 and P3, that is, directly at the termination of the intervention and one hour thereafter. A small positive effect on hemodynamics in early sepsis, during and in close relation to an intervention with HMW-HA is thus discernible in the present study.

Lactate increase was more pronounced in the intervention group as a function of time. This was not accompanied by a significant difference in pH or BE between the two groups (two-way ANOVA). However, comparing the two groups hourly, there was a tendency favoring the control group, with a statistically significant difference in BE, at P5 and P6. While there are several explanations to hyperlactatemia alone in sepsis, association to metabolic acidosis is most commonly interpreted as due to cardiovascular dysfunction or tissue hypoperfusion. Hypoxic hyperlactatemia can be explained by low cardiac output states and/or volume depletion [30]. In the present study, neither CO, SVV nor hemoconcentration differed between groups as a function of time. Oxygen extraction ratio and SvO_2_ also changed comparably in the two groups throughout the experiment. The difference in lactate can thus not be explained by a difference in any of these global measures of oxygen consumption/delivery mismatch. Hyperlactatemia, especially when refractory to resuscitation, is associated with increased mortality in sepsis [4,30,31]. In our fluid restrictive model of peritonitis, the finding of higher lactate in the intervention group is hampered by the short observation period. A more balanced resuscitation strategy might be of value to draw definite conclusions of potential benefit or harm.

Syndecan 1 and heparan sulfate in plasma are both sensitive markers of shedding of endothelial glycocalyx [32]. VAP 1 correlates with increased plasma concentrations of syndecan 1 in septic shock [33]. In our study, there was no difference in measured concentrations of syndecan 1, heparan sulfate or VAP 1 between groups. These findings suggest that HMW-HA alone does not exert a protective effect on the glycocalyx in early peritonitis sepsis. Norepinephrine requirements were also similar between the two groups as well as total protein, indicating a similar intravascular status in the two groups. Even though our model of postoperative peritonitis with source control is clinically relevant, an animal model can never fully replicate the human sepsis syndrome, due to possible differences in host response to both insult and intervention. Furthermore the fluid restrictive model enabled us to minimize the potential shedding of glycocalyx from crystalloid infusion per se, however it did not mimic the care of the septic patient, in which fluid resuscitation forms an integral part. Time control animals were stable throughout the experiment confirming the robustness of our model.

In conclusion, in this study, contrary to our hypothesis, HMW-HA infusion in a fluid restrictive model of early peritonitis sepsis did not prevent capillary leak and/or preserve intravascular volume. Whilst the infusion of HMW-HA was associated with higher CO and diastolic blood pressure, more pronounced hyperlactatemia with concomitant acidosis at the end of the experiment casts doubts on any potential beneficial effects of HMW-HA. For future studies, a balanced resuscitation strategy should be considered, with continuous infusion of HMW-HA, possibly with the addition of chondroitin sulphate (16, 21).

## Acknowledgements

The authors express their sincere gratitude to Kerstin Ahlgren, Liselotte Pihl, Mariette Andersson, and Maria Swälas for their assistance and support during the experiments at the Hedenstierna Laboratory of Uppsala University, Sweden.

## Notes

### Competing Interest Statement

The authors have declared no competing interest.

